# Assembly history modulates vertical root distribution in a grassland experiment

**DOI:** 10.1101/2021.08.24.457510

**Authors:** Inés M. Alonso-Crespo, Emanuela W.A. Weidlich, Vicky M. Temperton, Benjamin M. Delory

## Abstract

The order of arrival of plant species during assembly can affect the structure and functioning of grassland communities. These so-called priority effects have been extensively studied aboveground, but we still do not know how they affect the vertical distribution of roots in the soil and the rooting depth of plant communities. To test this hypothesis, we manipulated the order of arrival of three plant functional groups (forbs, grasses and legumes) in a rhizobox experiment. Priority effects were created by sowing one functional group 10 days before the other two. Rhizoboxes in which all functional groups were sown simultaneously were used as controls. During the experiment, the mean rooting depth of plant communities was monitored using image analysis and a new methodological approach using deep learning (RootPainter) for root segmentation. At harvest, we measured aboveground (community and species level) and belowground (community level) biomass, and assessed the vertical distribution of the root biomass in different soil layers. At the community level, all scenarios where one functional group was sown before the other two had similar shoot and root productivity. At the species level, two forbs (*Achillea millefolium* and *Centaurea jacea*) benefited from arriving early, and one legume (*Trifolium pratense*) had a disadvantage when it was sown after the grasses. Priority effect treatments also affected the vertical distribution of roots. When grasses were sown first, plant communities rooted more shallowly than when forbs or legumes were sown first,. In addition, roots moved down the soil profile 24% more slowly in grasses-first communities. Our results highlight that plant functional group order of arrival in grassland communities can affect the vertical distribution of roots in the soil and this may have implications for species coexistence.

## Introduction

**T**he order and timing of past biotic (e.g., species immigration) and abiotic (e.g., disturbances) events during community assembly can affect the way species interact with one another in ecological communities, making them historically contingent (Chase, 2003; Fukami, 2015). This historical contingency is often caused by priority effects (Drake, 1991). Typically, priority effects arise when species establishing early during assembly affect the persistence and growth of later arrivers (Kardol et al., 2013; Vannette and Fukami, 2014; Werner and Kiers, 2015; Delory et al., 2019; Temperton et al., 2016). If priority effects are strong enough to influence species coexistence, they can strongly affect the structure and functioning of ecological communities (Ejrnaes et al., 2006; Fukami et al., 2010; Martin and Wilsey, 2012; Sarneel et al., 2016; Werner et al., 2016; Weidlich et al., 2017). In fact, alternative stable states (Sutherland, 1974), alternative transient states (Fukami and Nakajima, 2011), and compositional cycles (Steiner and Leibold, 2004) are all possible long-term consequences of priority effects (Fukami, 2015).

In grasslands, priority effects have been studied mainly as a tool for ecological restoration to manipulate interactions between native and non-native species or as a way to foster ecosystem multifunctionality (Young et al., 2017; Hess et al., 2019; Weidlich et al., 2021). So far, most of the controlled and field experiments that have manipulated the sequence of arrival of species or plant functional groups (PFGs) have found strong aboveground priority effects on species diversity and aboveground productivity (Martin and Wilsey, 2012; von Gillhaussen et al., 2014; Weidlich et al., 2017), with implications for the resistance of plant communities to invasion (Hess et al., 2020). Perhaps surprisingly, much less is known about plant order of arrival effects on the productivity and distribution of roots in different soil layers, at both the community and species level.

The overall productivity of roots and their distribution in the soil are the result of two main factors: (1) the anatomical, morphological and architectural characteristics of the species populating the community (e.g., root growth form, root traits, etc.), and (2) the phenotypic plasticity of the roots of individual species in response to the biotic (e.g., neighbour identity and density, species richness, etc.) and abiotic (e.g., resource availability, soil texture, etc.) environment (Herben et al., 2018; Bakker et al., 2019; Chen et al., 2020; Case et al., 2020; Lepik et al., 2021). To date, much of what we know about the biotic and abiotic factors affecting the production and distribution of root biomass in grassland soils comes from experiments that did not manipulate the timing and/or order of plant species arrival. For instance, the effect of plant species and functional group richness on root dynamics and distribution at the community and species level has been a very active area of research in ecology (de Kroon et al., 2012). Biodiversity – ecosystem functioning (BEF) experiments that manipulated plant diversity without manipulating plant arrival order often found that root biomass production increases with plant species richness (Mommer et al., 2010; Mueller et al., 2013; Ravenek et al., 2014; Oram et al., 2018; Jesch et al., 2018; Zeng et al., 2021). However, the effect of plant diversity on the vertical distribution of roots has not been as clear, with most of the roots accumulating at the top of the soil (Mommer et al., 2010; de Kroon et al., 2012; Luo et al., 2021). Recently, two meta-analyses showed that increasing plant species richness does not seem to affect the rooting depth of plant communities (Barry et al., 2020; Peng and Chen, 2021). With the exception of one study that reported that high diversity plots in a grassland field experiment had a higher proportion of root biomass allocated to deeper soil layers (Mueller et al., 2013), there is little evidence supporting an increase in vertical root niche differentiation with increased plant diversity in either grasslands (Mommer et al., 2010; Ravenek et al., 2014; Oram et al., 2018) or forests (Valverde-Barrantes et al., 2015; Luo et al., 2021; Zeng et al., 2021). Differences in rooting patterns between plant functional groups also received mixed support, with some studies reporting that grasses rooted more superficially than forbs (Bakker et al., 2019; Chen et al., 2020), while others found no difference between forbs and grasses (Mommer et al., 2010; Ravenek et al., 2014; Oram et al., 2018).

If species were to arrive sequentially, and not at the same time as in most BEF experiments, one can expect the productivity and distribution of roots to be affected by plant order of arrival. For instance, if species A arrives before species B at a site, species A will start taking up resources earlier and will reduce the availability of essential soil resources for species B (niche pre-emption) (Kardol et al., 2013). In addition, species A might create soil legacies (niche modification) that will affect the establishment of species B (Kardol et al., 2007; Grman and Suding, 2010; Delory et al., 2021). Both niche pre-emption and niche modification mechanisms could trigger plastic root responses in some later arriving species, which could affect how they develop and position their roots in the soil (Chen et al., 2020). These species-specific root responses could then spread to the community level and affect the overall root distribution and average rooting depth of plant communities. So far, both controlled and field experiments have revealed that manipulating the order of arrival of PFGs can affect root productivity in the topsoil of mesic grasslands, with lower standing root biomass and root length density when leguminous species were sown first (Körner et al., 2008; Weidlich et al., 2018b). Whether this pattern was due to changes in total root productivity, vertical root distribution, or both is still unknown.

In this paper, we present the results of a rhizobox experiment designed to test how PFG order of arrival (forbs, grasses and legumes) in mesic grasslands affects root productivity and the vertical distribution of roots at the early stages of plant community assembly. Our experiment tested the following hypotheses:

1. The above- and belowground productivity of plant communities varies depending on whether forbs, grasses, or legumes were sown first;
2. The vertical distribution of roots depends on PFG order of arrival during assembly, with communities in which forbs or legumes were sown first rooting deeper than communities in which grasses were sown first.

## Materials and Methods

### Experimental design and setup

We conducted a rhizobox experiment at a greenhouse located in Lüneburg, Germany (53°14’23.8”N, 10°24’45.5”E) in August-September 2017. The main goal of this experiment was to test the influence of assembly history on the vertical root distribution of a plant community typical of mesic grasslands. We manipulated assembly history by altering the order of arrival of three plant functional groups (PFGs): non N_2_-fixing forbs (forbs), N_2_-fixing forbs (legumes), and grasses. Each plant community consisted of nine species: three forbs (*Achillea millefolium, Leucanthemum vulgare, Centaurea jacea*), three legumes (*Lotus corniculatus, Medicago sativa, Trifolium pratense*), and three grasses (*Dactylis glomerata, Festuca rubra, Holcus lanatus*).

One week before starting the experiment, 35 rhizoboxes (58×26.6×2 cm) were filled with a 5 mm-sieved mixture of sand (30%, v/v) and potting soil (70%, v/v). Each rhizobox was then watered with 100 ml of tap water and randomly assigned to an experimental treatment. Four days later, all rhizoboxes received again 50 ml of tap water.

In this experiment, we manipulated assembly history using five different PFG order of arrival scenarios: synchronous 1 (Sync1; all PFGs were sown simultaneously at the first sowing event), synchronous 2 (Sync2; all PFGs were sown simultaneously at the second sowing event), forbs-first (F-first; forbs sown before grasses and legumes), grasses-first (G-first; grasses sown before forbs and legumes), and legumes-first (L-first; legumes sown before forbs and grasses). Each PFG order of arrival scenario was replicated 7 times. Note that we set up two synchronous treatments so that the performance of plants that were sown at the first or second sowing event could be compared directly to plants that had grown for the same length of time. The time interval between the first and second sowing events was 10 days. The position of the functional groups inside the rhizoboxes was fixed (Fig. 1), but the position of each species within a functional group was assigned randomly. In each rhizobox, all plant individuals were equidistantly spaced (2.7 cm). A few days after each sowing event, the seeds that did not germinate were replaced by seedlings of the same species that had been allowed to germinate in Petri dishes filled with the same soil as the one used in the rhizoboxes. The Petri dishes were stored vertically in a plastic tray next to the experiment. Each rhizobox was watered regularly with 20 to 50 ml of tap water. Over the entire duration of the experiment, each rhizobox received a total of 770 ml of tap water. All rhizoboxes were placed in plastic containers (5 rhizoboxes per container) and were inclined at a 45° angle relative to the vertical in order to allow the roots to grow along the transparent front window. In each container, the front window of the first rhizobox was covered with a black plastic plate to prevent root exposition to light, and the back of the last rhizobox was covered with a white polystyrene plate to avoid overheating. The position of the rhizoboxes inside the greenhouse was regularly randomized during the experiment.

**Fig. 1.**
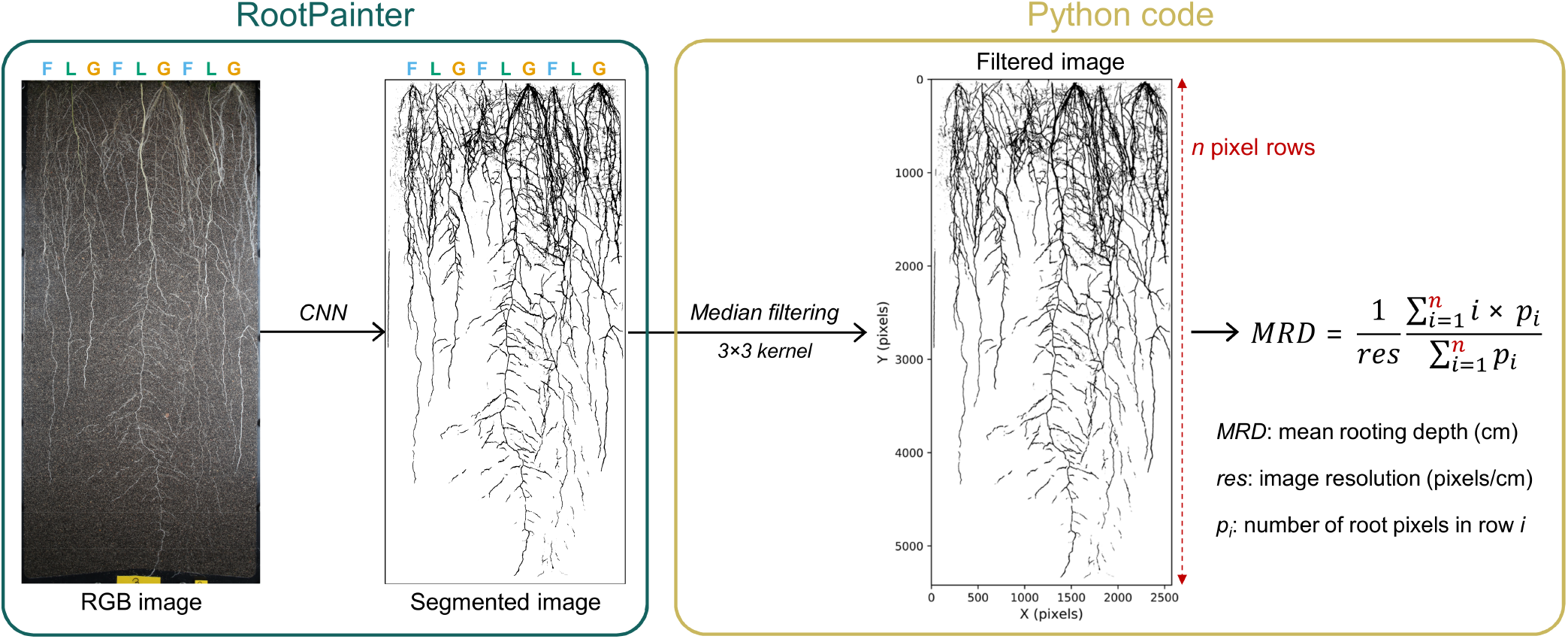
Description of the image analysis pipeline used for non-destructively estimating the mean rooting depth (MRD) of plant communities growing in rhizoboxes. Our approach included three main steps: (1) training a convolutional neural network (CNN) using RootPainter to detect plant roots in our images, (2) segmenting all images taken during the experiment with the best performing model, and (3) estimating the average rooting depth of each plant community at each time point using a Python procedure run in a Jupyter Notebook. F, forb species; G, grass species; L, legume species.

### Root image acquisition

Root images were acquired three times a week (Monday, Wednesday, Friday) using a digital camera (Canon EOS 5D Mark III) equipped with a 28 mm lens (Canon EF 28mm f/2.8) and connected to a computer (Delory et al., 2018). In addition to the camera and its associated holder, our image acquisition system consists of a metallic frame holding the rhizobox vertically and two LED tubes (4300 K, 60 cm length) positioned laterally (raking lighting) to provide uniform lighting conditions over the entire height of a rhizobox. The camera, the camera holder, the LED tubes, and the frame holding the rhizobox are installed inside a closed box whose internal walls are entirely covered with dark fabric. Images were acquired using our camera’s remote live view and shooting option before being saved on a computer using both compressed (jpeg) and uncompressed (raw CR2) file formats.

### Harvest and measurements

Forty-four days after the start of the experiment (first sowing event), plant shoots were harvested and stored separately for each species. Shoot samples were dried at 60°C until constant mass was reached, and weighed.

At harvest, rhizoboxes were opened by carefully removing the transparent front window. Using a sharp knife, the soil profile was divided into six 10-cm layers (0-10, 10-20, 20-30, 30-40, 40-50, >50 cm). For each soil layer, roots were extracted from the soil under running water and stored at -20°C until further processing. Fine root washing was performed in the lab by carefully removing soil particles adhering to the roots following Delory et al. (2018). Clean root samples were then dried at 60°C for at least 48 h, and weighed. For each rhizobox, total root productivity was calculated by summing the root dry weight values measured in all soil layers.

### Analysis of the vertical distribution of roots in the rhi-zoboxes

Two strategies were used to investigate the effect of PFG order of arrival on vertical root distribution: (1) modelling root biomass distribution in the soil (i.e., root dry weight as a function of depth), and (2) modelling the temporal evolution of the average rooting depth of plant communities (i.e., rooting depth as a function of time).

Root biomass distribution in the soil was modelled using a zero-altered gamma model (also referred to as a hurdle model) following Zuur and Ieno (2016). This choice of statistical model was motivated by the fact that our dataset consisted of zero-inflated continuous data (Y*≥* 0) since some layers did not contain any roots at harvest. The relationship between root biomass production and soil depth was investigated using a hurdle model combining a binomial generalized linear mixed-effect model (binomial GLMM) and a gamma generalized linear mixed-effect model (gamma GLMM). The binomial GLMM component was used to model the relationship between the presence/absence of roots (Y=1 or 0) and soil depth, while the gamma GLMM part modelled the relationship between non-zero root biomass data (Y>0) and soil depth. GLMMs were fitted using PFG order of arrival (5 levels), soil depth (continuous variable), and their interaction as fixed effects. To account for the fact that root biomass values measured at different depths in the same rhizobox were not independent, rhizobox ID was used as a random effect in the models. The binomial GLMM was fitted using a logit link function, while the gamma GLMM was fitted using a log link function. GLMM models were fitted using the MASS package (Venables and Ripley, 2002).

The mean rooting depth (MRD) of plant communities was estimated using image analysis as the depth value above which 50% of the roots are located (Mommer et al., 2010; Freschet et al., 2020). For each image taken during the experiment, a value of MRD was estimated using the approach illustrated in Fig. 1. First, root images were cropped in ImageJ using a custom macro to select only the zone of soil containing roots (Schindelin et al., 2012). These cropped images were used to create a training dataset of 600 smaller images (height: 612 to 990 pixels; width: 643 to 872 pixels). This training dataset was created with RootPainter in two steps: (1) randomly selecting 300 images from the 665 images taken during the experiment (35 rhizoboxes × 19 time points), and (2) randomly selecting two subregions of each image selected in step 1 (Smith et al., 2020). Then, RootPainter was used to annotate a number of training images and train a convolutional neural network (CNN) to detect roots in our images (Smith et al., 2020). Once model predictions successfully identified most of the roots present in our training images, the training was stopped and the best performing model was used to segment the cropped version of the 665 images taken during the experiment. All segmented images were then automatically processed using a custom Python procedure consisting of the following steps: (1) loading a segmented image and converting it to a binary format where a root pixel has a value of one and a background pixel has a value of zero, (2) applying a median filter (3×3 kernel) to remove noise, and (3) calculating a value for MRD using the equation shown in Fig. 1. For each image, MRD was calculated as the sum of the total number of root pixels in row *i* multiplied by the depth of row *i* divided by the total number of root pixels in that image. MRD values calculated by image analysis and MRD values calculated with root biomass data using the same approach were highly positively correlated (see Figure 5a). The temporal evolution of MRD was modelled using a linear mixed-effect model. The fixed component of the model contained PFG order of arrival (5 levels), the number of days after sowing each rhizobox (continuous variable), and their interaction. Considering that the rooting depth in each rhizobox was measured at multiple time points, the temporal evolution of MRD was modelled using a random slope and a random intercept for each rhizobox. The linear mixed-effect model was fitted using the lme4 R package (Bates et al., 2015).

### Data analysis

As plant shoots were harvested separately for each species, shoot biomass data was analysed at both species and community levels. Root biomass data, however, was only analysed at the community level (roots from different species could not be separated from each other). When analysing and interpreting the shoot and root biomass data presented in this paper, we considered recent calls to stop using P-values in a dichotomous way and stop declarations of “statistical significance” (Amrhein et al., 2019; Wasserstein et al., 2019). To do so, we reported effect sizes (difference between treatment means) and their 95% confidence intervals computed by bootstrap resampling (20,000 iterations). Following Amrhein et al. (2019), 95% confidence intervals will be referred to as compatibility intervals in the remainder of the paper. Since manipulating PFG order of arrival implies that seeds of different species were not sown at the same time in all rhizoboxes, meaningful comparisons have to be defined a priori so that the calculated effect sizes reflect plant order of arrival effects, and not differences in plant age. When shoot biomass data collected at the species level was analysed, care was taken to compare plant individuals that grew for the same length of time. This means that the productivity of a species in a priority effect treatment (F-first, G-first, or L-first) was compared either to the productivity achieved by the same species in the Sync1 treatment (if the target species was sown at the first sowing event), or Sync2 treatment (if the target species was sown at the second sowing event). For shoot and root productivity data measured at the community level, effect sizes were calculated by comparing priority effect treatments to each other, but not to synchronous treatments. For each response variable, effect sizes were assessed by comparing the mean values and their compatibility intervals.

Data analysis was performed in R 4.0.5 (R Core Team, 2021). Plots were created using the R packages ggplot2 (Wickham, 2016), ggeffects (Lüdecke, 2018) and ggpubr (Kassambara, 2020).

## Results

### Priority effects on shoot and root productivity

At the community level, we found that the total shoot productivity of F-first, G-first, and L-first communities did not markedly differ from each other (Fig. 2a). Although F-first communities were on average 15% and 17% more productive than G-first and L-first communities, respectively, our data are also compatible with an effect size of zero (i.e., no difference in total shoot productivity). Similarly, we did not find any difference in total root pro-ductivity between F-first, G-first, and L-first communities (Fig. 2b). As expected, the total aboveground and below-ground productivity was highest in the Sync1 treatment, but lowest in the Sync2 treatment (Fig. 2a-b).

**Fig. 2.**
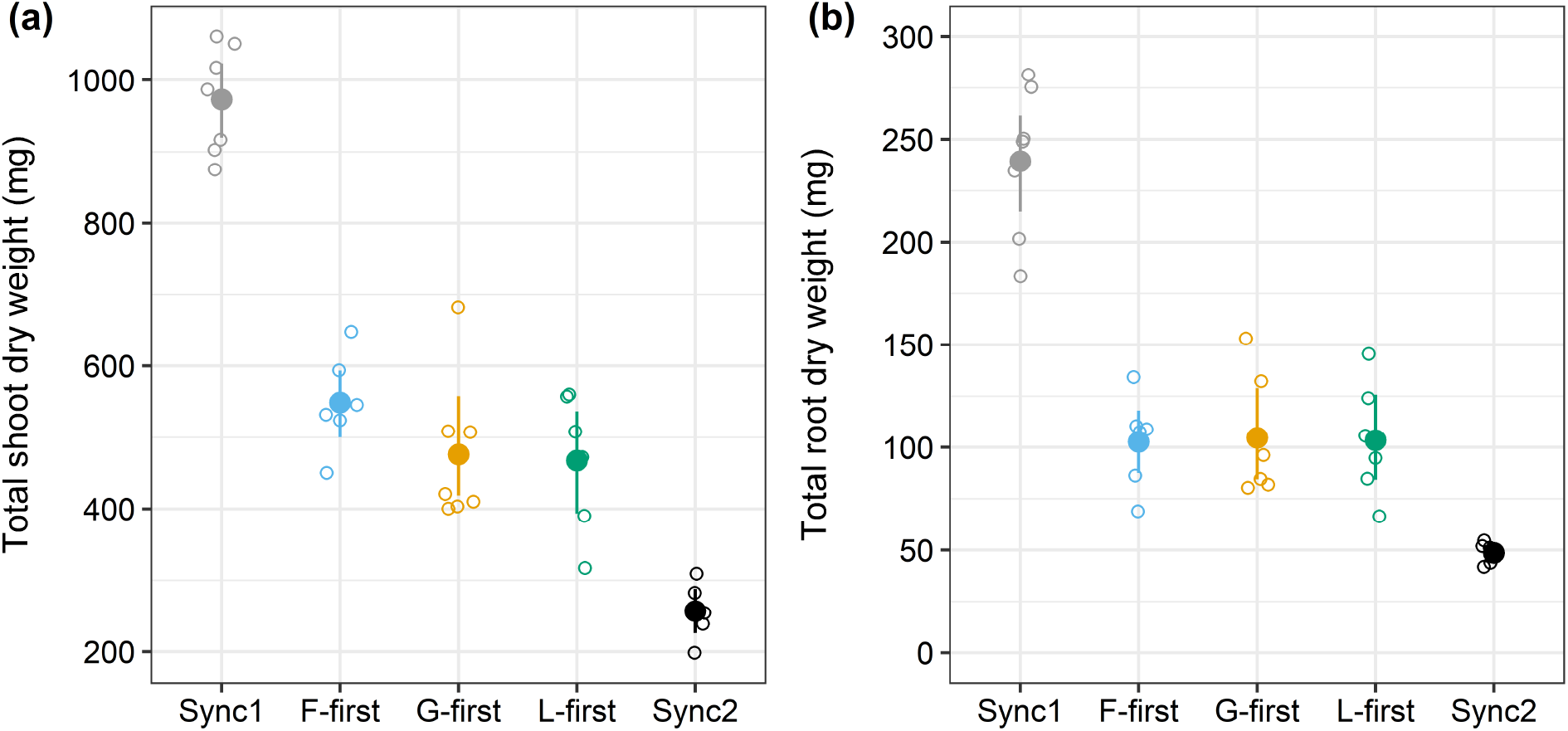
Effect of PFG order of arrival on the total shoot (a) and root (b) productivity of plant communities. For each treatment, mean values and compatibility intervals computed using non-parametric bootstrap are shown (n=5-7). Individual observations are displayed as empty dots. Sync1, all PFGs sown at the same time at the first sowing event; F-first, forbs sown 10 days before grasses and legumes; G-first, grasses sown 10 days before forbs and legumes; L-first, legumes sown 10 days before forbs and grasses; Sync2, all PFGs sown at the same time at the second sowing event.

At the species level, two forb species clearly benefited from arriving early. Both *Achillea millefolium* (+87%) and *Centaurea jacea* (+67%) were more productive above-ground in the F-first treatment than in the Sync1 treatment (Fig. 3). Results also showed that one legume species, *Trifolium pratense*, suffered from arriving after the grasses. In the G-first scenario, *T. pratense* had a lower shoot biomass productivity than in the Sync2 (−34%) and F-first (−24%) treatments.

**Fig. 3.**
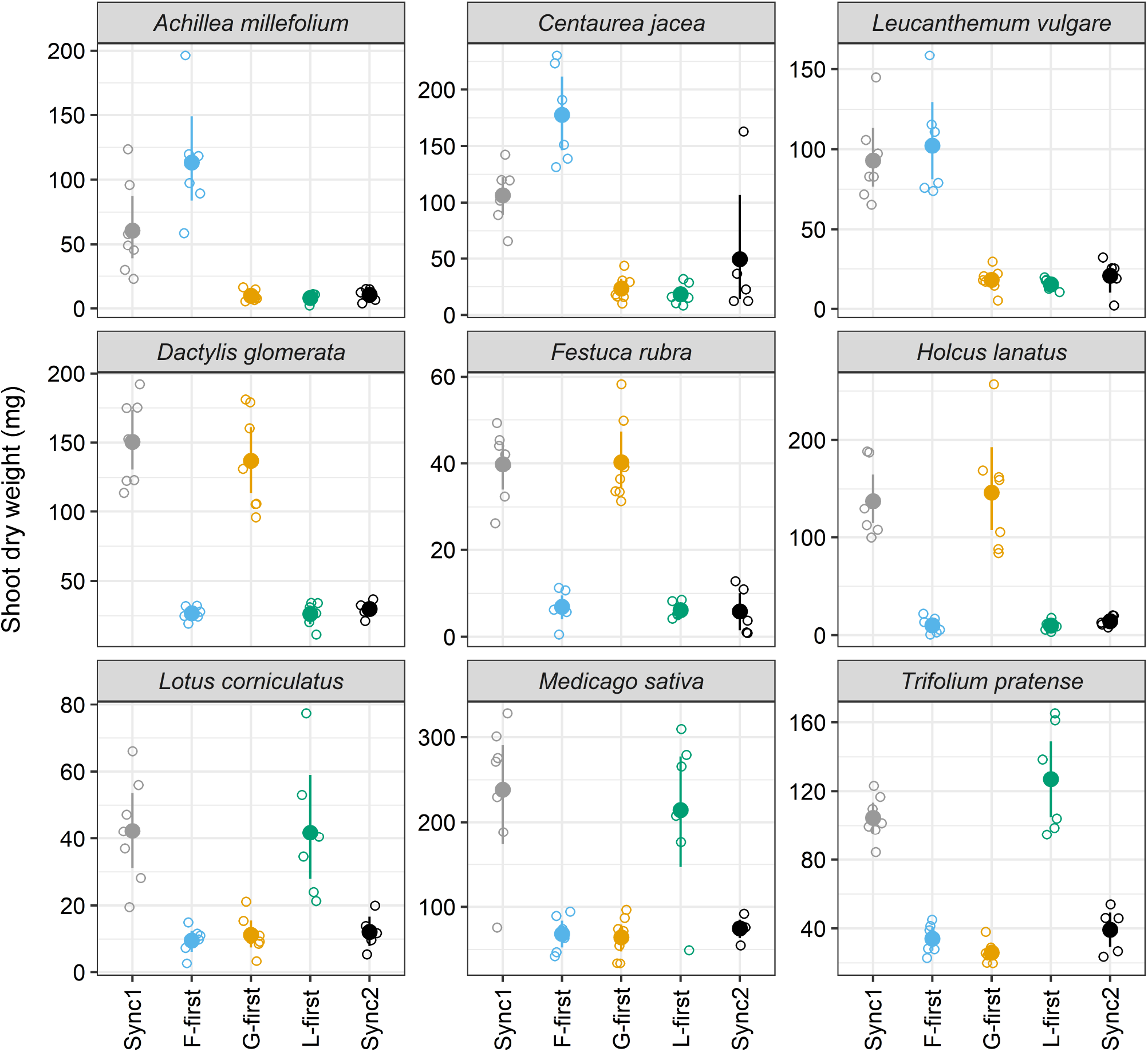
Effect of PFG order of arrival on the shoot productivity of individual species. For each treatment, mean values and compatibility intervals computed using non-parametric bootstrap are shown (n=5-7). Individual observations are displayed as empty dots. Sync1, all PFGs sown at the same time at the first sowing event; F-first, forbs sown 10 days before grasses and legumes; G-first, grasses sown 10 days before forbs and legumes; L-first, legumes sown 10 days before forbs and grasses; Sync2, all PFGs sown at the same time at the second sowing event.

### Root biomass distribution in the soil

Despite the fact that we did not observe any difference in total root productivity between the F-first, G-first, and L-first scenarios, we found that the vertical distribution of root biomass in the soil was strongly affected by PFG order of arrival (Fig. 4). When grasses were sown first (G-first), a larger proportion of the root biomass was found at the top of the soil, and the observed decrease in root productivity with depth was stronger than when forbs or legumes were sown first (Fig. 4). In fact, we did not find any roots located deeper than 50 cm when grasses were sown first. This was not the case when forbs or legumes were sown first since roots located deeper than 50 cm were mostly present (all but one rhizobox) (Fig. S1).

**Fig. 4.**
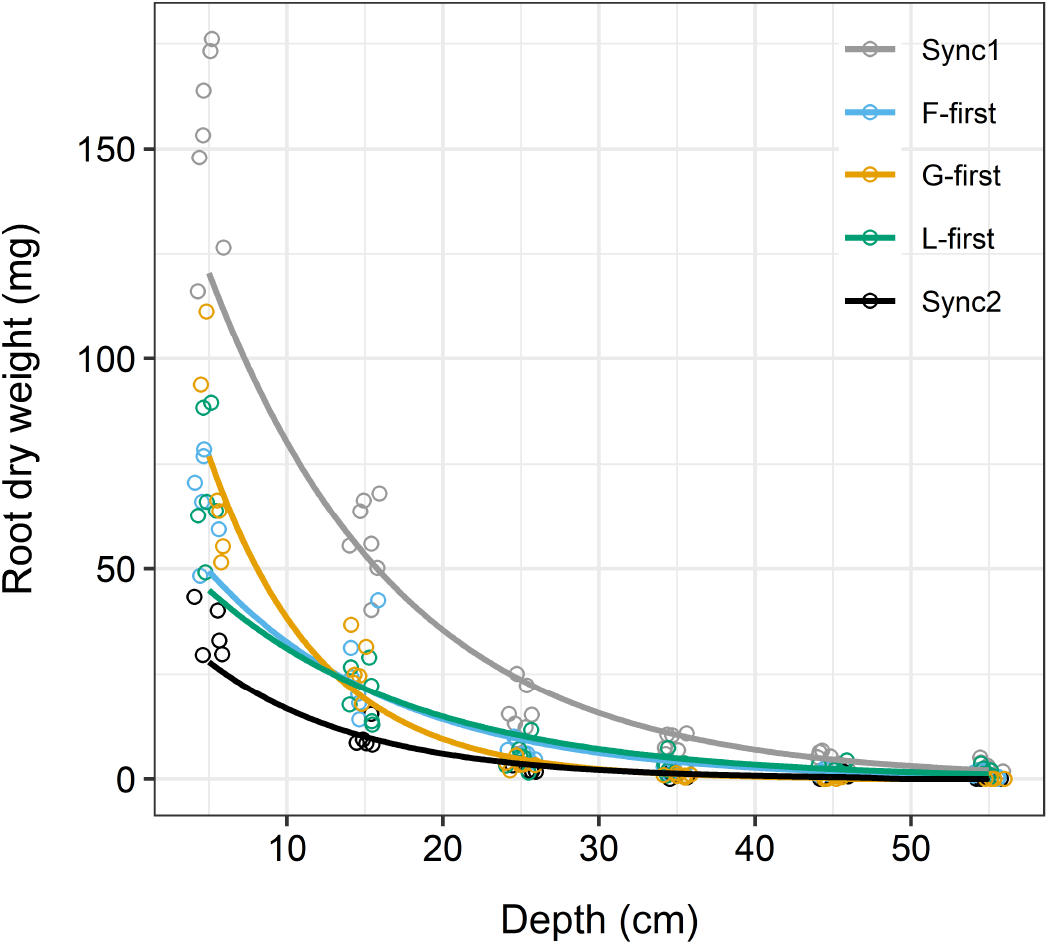
Vertical root biomass distribution in the soil is affected by PFG order of arrival. Lines represent ZAG model fits for each PFG order of arrival scenario. Root biomass production values are shown for each soil layer. Individual observations are displayed as empty dots. Sync1, all PFGs sown at the same time at the first sowing event; F-first, forbs sown 10 days before grasses and legumes; G-first, grasses sown 10 days before forbs and legumes; L-first, legumes sown 10 days before forbs and grasses; Sync2, all PFGs sown at the same time at the second sowing event.

### Temporal evolution of the rooting depth of plant communities

Our results showed that the temporal evolution of the mean rooting depth (MRD) of plant communities was dependent on PFG order of arrival (Fig. 5b-c). In particular, we found that G-first communities rooted more shallowly than the others. Roots moved down the soil profile more slowly for G-first communities. On average, MRD increased by 2.7 mm day-1 in synchronous (Sync1 and Sync2), F-first and L-first communities, whereas it increased by only 2.1 mm day-1 in G-first communities, which represents a 24% decrease in progression rate. At the last observation date (day 44), Sync1, F-first and L-first communities rooted between 3.1 and 4.6 cm deeper than G-first communities (Fig. 5b-c).

**Fig. 5.**
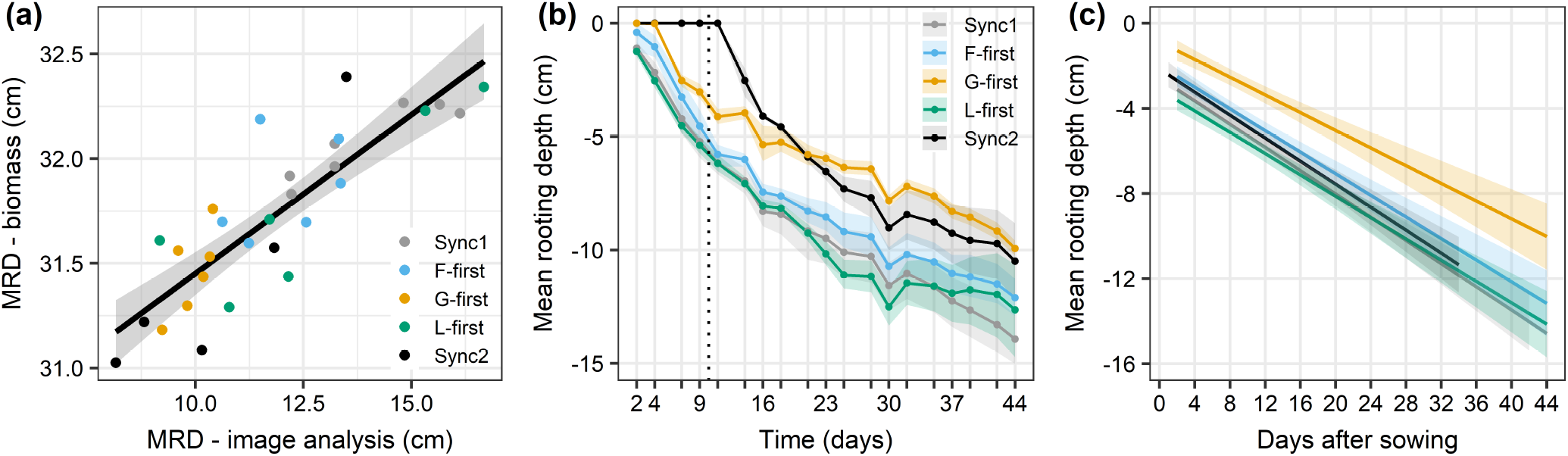
The temporal evolution of rooting depth is affected by PFG order of arrival. (a) Relationship between MRD values computed using root biomass data and MRD values computed using image analysis (Fig. 1); (b) Temporal evolution of the mean rooting depth (MRD – image analysis) of plant communities for each PFG order of arrival scenario; (c) Linear mixed-effect model fits describing the temporal evolution of MRD for each plant community type. In panel b, average MRD values were represented at each time point as filled dots. Sync1, all PFGs sown at the same time at the first sowing event; F-first, forbs sown 10 days before grasses and legumes; G-first, grasses sown 10 days before forbs and legumes; L-first, legumes sown 10 days before forbs and grasses; Sync2, all PFGs sown at the same time at the second sowing event.

## Discussion

### Species-specific effects of PFG order of arrival on shoot productivity

At the community level, we found that the total shoot productivity was similar if forbs, grasses, or legumes were sown first. As expected, the yield of communities in which one PFG was sown before the other two was intermediate between the yields of synchronous communities (Sync1 and Sync2). These results can appear surprising because they contradict findings from other controlled experiments that manipulated PFG order of arrival. Indeed, both von Gillhaussen et al. (2014) and Körner et al. (2008) found that grassland plant communities in which legumes were sown a few weeks before grasses and forbs were amongst the most productive. In a field experiment that manipulated PFG order of arrival in a mesic grassland, Weidlich et al. (2017) also found a similar pattern, but results were quite variable from year to year. We suggest that the shorter sowing interval between early and late arrivers used in this study (10 days) as well as the shorter duration of our experiment (6 weeks), which were imposed by our experimental set up (i.e., plants grown in rhizoboxes), are likely to be the main factors explaining why our study yielded unusual results regarding how PFG order of arrival affected the productivity of plant communities.

In comparison with a scenario where all species were sown at the same time (Sync1 and Sync2), only three species performed differently when they arrived earlier or later than other species in the community. While the growth of *T. pratense* was only negatively affected when grasses were sown first (i.e., negative priority effect), two of the three forb species used in this experiment clearly benefited from arriving early in the community. Both *C. jacea* and *A. millefolium* had a greater shoot productivity when forbs were sown first (F-first > Sync1). The forb *L. vulgare*, however, reached the same productivity level when it arrived earlier (F-first) or at the same time as the other species (Sync1). Although the exact mechanisms (niche pre-emption vs niche modification sensu Fukami (2015)) that created negative priority effects for *T. pratense* when grasses were sown first are still unknown, a possible explanation for the fact that *C. jacea* and *A. millefolium* benefited from arriving early (while *L. vulgare* did not) might be related to their different rooting strategies. In a common garden experiment, Bakker et al. (2019) found that *C. jacea* and *A. millefolium* had a greater deep root fraction and a lower specific root length than *L. vulgare*. In comparison with *C. jacea* and *A. millefolium, L. vulgare* seems to have another rooting strategy that aims to explore the topsoil more efficiently by producing longer and finer roots and invest less biomass in deeper soil layers (Bakker et al., 2019). Considering that *C. jacea* and *A. millefolium* grew better when they were sown before grasses and legumes, our results suggest that root traits such as deep root fraction may be important in mediating the susceptibility of plant species to plant order of arrival, with deep-rooting species benefiting more from arriving early than shallow-rooting species. This hypothesis, however, still requires further investigation.

### Total root productivity was similar when forbs, grasses or legumes were sown first, but sowing grasses before forbs and legumes led to more shallow-rooted communities

As with the aboveground productivity results, total root biomass production was similar when forbs, grasses or legumes were sown first. The total root productivity achieved in communities where one PFG was sown earlier was intermediate between root productivity values measured in synchronous communities (Sync1 and Sync2). These results do not agree with our hypothesis that belowground productivity of plant communities would vary depending on whether forbs, grasses, or legumes were sown first. This is unexpected considering that previous studies found that grassland plant communities are less productive belowground when legumes are given a head start (Körner et al., 2008; Weidlich et al., 2018b). As with the aboveground productivity results, the constraints imposed by our experimental set up are probably one of the main reasons for these differences.

Although we did not find any difference in total root productivity when forbs, grasses or legumes were sown first, we found that manipulating PFG order of arrival had a strong effect on the vertical distribution of roots in the soil. In particular, our results showed that communities in which forbs or legumes were sown first rooted deeper than communities in which grasses arrived a few days before the other PFGs, thus confirming our second hypothesis. Root biomass distribution modelling showed that grasses-first communities had a greater proportion of roots allocated in upper soil layers. In addition, we found that roots progressed more slowly through the soil profile when grasses were sown before forbs and legumes. Remarkably, the temporal evolution of the average rooting depth was very similar between F-first, L-first and synchronous communities (Sync1 and Sync2). These results confirm that assembly history can modulate the vertical distribution of roots in grassland ecosystems, which is remarkable given the short time interval between the arrival of early and late species in our experiment (10 days). However, the underlying mechanisms that led to these effects are still unknown and further research is needed to determine the extent to which niche preemption and niche modification by early-arriving species played a role (Fukami, 2015).

Although strong differences in root distribution exist between individual plant species (Herben et al., 2018; Lepik et al., 2021), evidence for differences in root distribution between plant functional groups in grassland ecosystems has been mixed, with some studies reporting no difference in rooting depth between functional groups (Mommer et al., 2010; Ravenek et al., 2014; Oram et al., 2018), while others have indicated that forb species root deeper on average than grasses (Bakker et al., 2019; Chen et al., 2020). Since our plant communities had exactly the same species and functional group composition and only differed by the order of arrival of forbs, grasses and legumes, differences in rooting depth between species or functional groups is unlikely to be the only explanation behind our results. Instead, we believe that plastic and species-specific root responses to plant order of arrival are more likely to explain why grasses-first communities rooted more shallowly than the others did. Such plastic root responses affecting root allocation and foraging have been well documented in the past (Mahall and Callaway, 1991; Semchenko et al., 2007; Mommer et al., 2012; Kumar et al., 2020; Lepik et al., 2021) but, to date, there has been little evidence that the order of arrival of plants can affect the root distribution of individual species (Weidlich et al., 2018a). Studying the extent to which the behaviour of the roots of species inhabiting plant communities changes as a function of the order of arrival of plants, as well as studying how these plastic root responses would be reflected at the community level, requires information on the distribution of roots at the species level, which unfortunately was not available in our study. Given the above, as well the inherent limitations of any rhizobox experiment, we see two main avenues for future research aimed at better understanding the roles played by priority effects in root dynamics and their consequences for species coexistence: (1) non-destructively monitoring root development at different soil depths in the field using minirhizotrons (Rewald and Ephrath, 2013; Freschet et al., 2020), and (2) quantifying species relative abundance in root samples taken from plant communities at different soil depths using state-of-the-art molecular techniques (Wagemaker et al., 2021). Only by going underground can we improve our mechanistic understanding of priority effects in plant communities and their implications for species coexistence.

## Conclusions

Using root biomass distribution data and deep learning-based image analysis, we provided evidence that assembly history, and more particularly plant order of arrival during assembly, can modulate the vertical distribution of roots in a controlled grassland experiment. When grasses were sown before forbs and legumes, plant communities had the shallowest root distribution. This result was explained by the fact that (1) a greater proportion of root biomass was present at the top of the soil, and (2) roots progressed more slowly through the soil in communities in which grasses were sown first. Further research is needed to better understand how priority effects alter root dynamics at the community and species level, and how this may affect species coexistence. Field experiments that use a larger sowing interval between plant functional groups would be ideal for such an endeavour.

## Acknowledgements

The authors would like to thank Dr Thomas Niemeyer (Leuphana University of Lüneburg, Germany) for his outstanding technical support and Abraham George Smith (University of Copenhagen, Denmark) for his extensive advice that enabled us to successfully use RootPainter to analyse our root images.

## Data availability

The raw data, the RootPainter model used to segment root images (.pkl), and the annotated R and Python codes supporting the findings of this study are available in Zenodo.

---

**Fig. S1.**
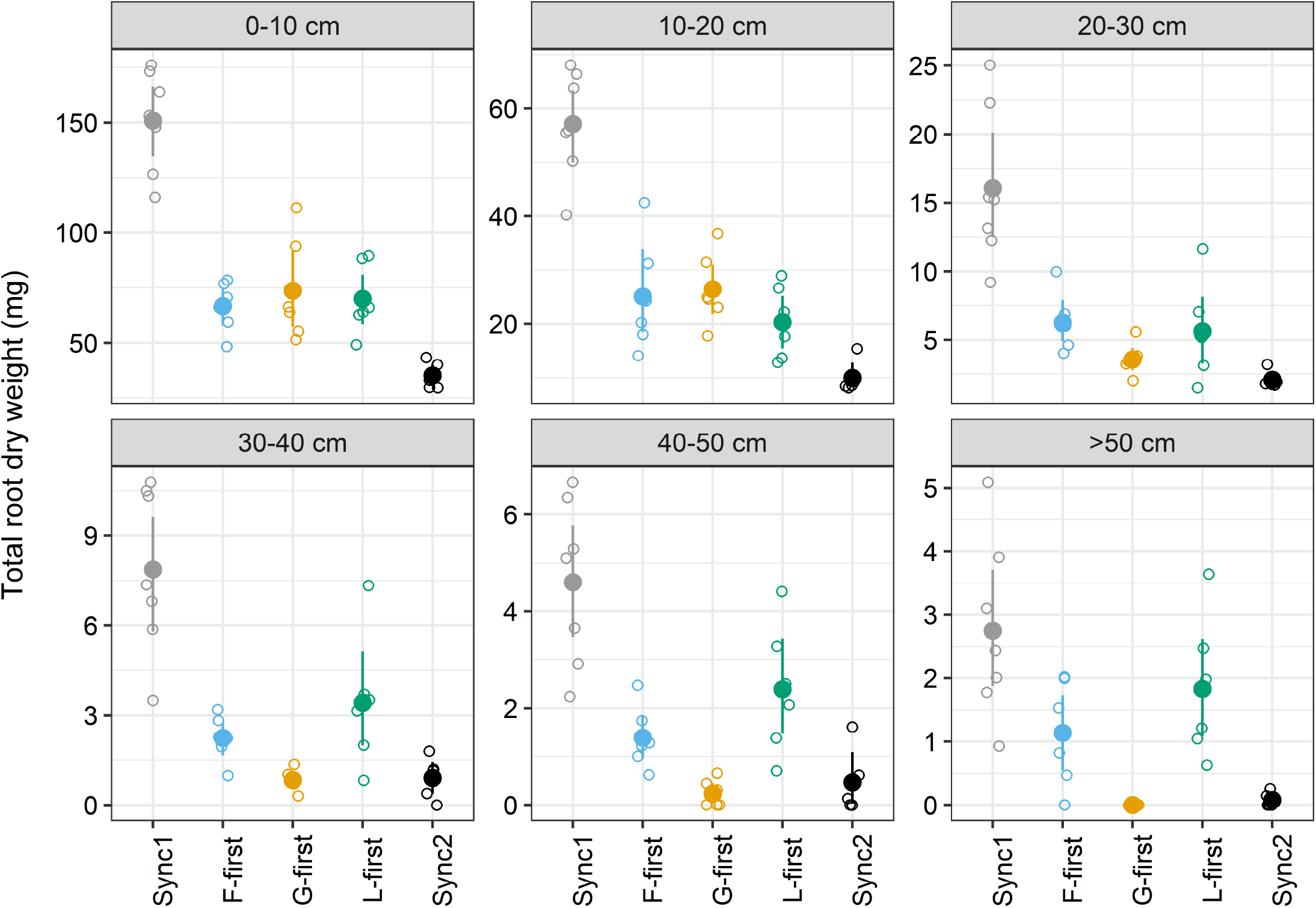
Root biomass distribution in different soil layers. For each treatment, mean values and compatibility intervals computed using non-parametric bootstrap are shown (n=5-7). Individual observations are displayed as empty dots. Sync1, all PFGs sown at the same time at the first sowing event; F-first, forbs sown 10 days before grasses and legumes; G-first, grasses sown 10 days before forbs and legumes; L-first, legumes sown 10 days before forbs and grasses; Sync2, all PFGs sown at the same time at the second sowing event.

